# Previous SARS-CoV-2 infection increases B.1.1.7 cross-neutralization by vaccinated individuals

**DOI:** 10.1101/2021.03.05.433800

**Authors:** Benjamin Trinité, Edwards Pradenas, Silvia Marfil, Carla Rovirosa, Víctor Urrea, Ferran Tarrés-Freixas, Raquel Ortiz, Júlia Vergara-Alert, Joaquim Segalés, Victor Guallar, Rosalba Lepore, Nuria Izquierdo-Useros, Glòria Trujillo, Jaume Trapé, Carolina González-Fernández, Antonia Flor, Rafel Pérez-Vidal, Anna Chamorro, Roger Paredes, Ignacio Blanco, Eulalia Grau, Marta Massanella, Jorge Carrillo, Bonaventura Clotet, Julià Blanco

**Author notes:** **Corresponding author:** Julià Blanco, PhD, Senior Researcher, Institut de Recerca de la Sida. IrsiCaixa, IGTP, Hospital Germans Trias i Pujol, Ctra. de Canyet s/n. 2a Planta Maternal. 08916 Badalona. Barcelona. Equal contribution (Benjamin Trinité, Edwards Pradenas).

## Abstract

To assess the potential impact of predominant circulating SARS-CoV-2 variants on neutralizing activity of infected and/or vaccinated individuals, we analyzed neutralization of pseudoviruses expressing the spike of the original Wuhan strain, the D614G and B.1.1.7 variants. Our data show that parameters of natural infection (time from infection and infecting variant) determined cross-neutralization. Importantly, upon vaccination, previously infected individuals developed equivalent B.1.1.7 and Wuhan neutralizing responses. In contrast, uninfected vaccinees showed reduced neutralization against B.1.1.7.

**Funding:** This study was funded by Grifols, the *Departament de Salut* of the *Generalitat de Catalunya*, the Spanish Health Institute Carlos III, CERCA Programme/*Generalitat de Catalunya*, and the crowdfunding initiatives #joemcorono, BonPreu/Esclat and Correos.

---

The variant B.1.1.7 or 501Y.V1 has rapidly spread across Europe (1). Besides the D614G mutation, this variant contains 6 mutations and 3 deleted amino acids in the Spike protein. Major changes are the mutation N501Y in the Receptor Binding Domain (RBD); the deletion 69-70 which may increase transmissibility (2) and produces false negative in certain RT-PCR based diagnostic assays; and the mutation P681H, next to the furin cleavage site, that could impact antigenicity and enhance viral infectivity. Although there is no convincing evidence that B.1.1.7 causes more severe disease, several reports indicate that B.1.1.7 has a higher secondary attack rate making this viral variant 30-50% more transmissible (3). Importantly, this variant remains susceptible to some monoclonal and plasma antibodies from convalescent or vaccinated individuals (4–6).

To comparatively evaluate infectivity and neutralization sensitivity, we constructed HIV-1-based pseudoviruses expressing different variants of the S protein of SARS-CoV-2 as reported (7): the original sequence (Wuhan); the single point mutant D614G which developed during the first European outbreak and became ubiquitous at subsequent outbreaks; and the B.1.1.7 variant, that has recently become predominant in Europe (Figure 1A). The neutralization of plasma samples was assessed using pseudoviruses as previously described (8), calculating ID50 values as reciprocal plasma dilution. Pseudoviruses were generated in Expi293 cells by co-transfecting a defective reporter HIV and the corresponding Spike coding plasmid with a deletion of the last 19 amino acids (Supplementary Methods). Two days after transfection, pseudoviruses were harvested and titrated on ACE-2-HEK293T cells (Integral Molecular, USA). Consistent with previously reported data (9), viral evolution is associated with increased fifty-percent tissue culture infective dose (TCID50) *in vitro* for both the D614G and B.1.1.7 variants (Figure 1B). Spike expression in pseudovirus producing cells was confirmed by flow cytometry, showing comparable expression for all constructs (Supplementary Methods, Figure 1C).

**Figure 1.**
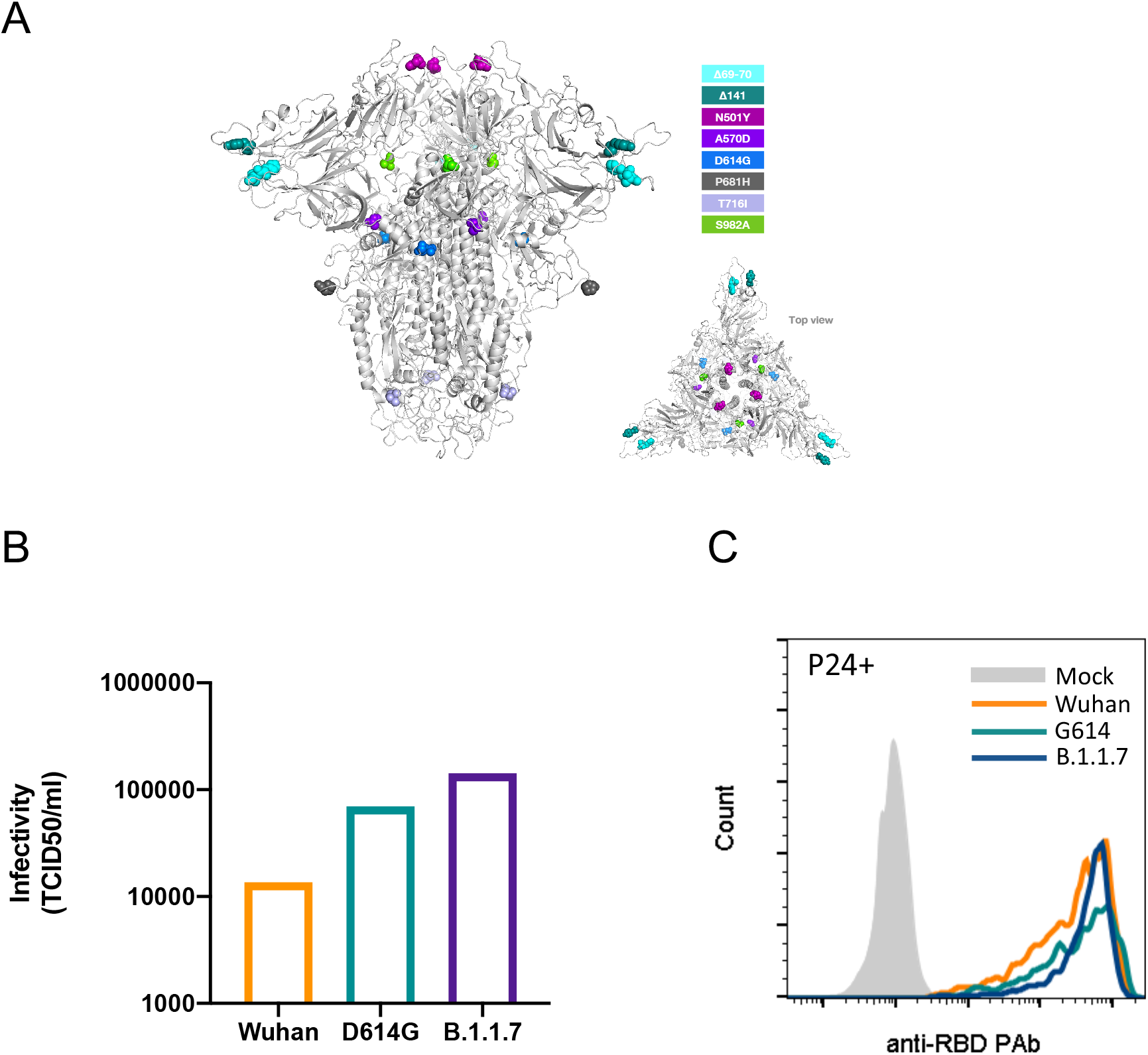
In vitro infectivity of SARS-CoV-2 variants. **A.** The different mutations identified in the B.1.1.7 variant are listed and their location in the Spike protein (side and top views) are shown. This variant also includes the D614G mutation. **B**. Infectivity of pseudoviruses recovered from Expi293 and titrated in human ACE-2 expressing 293T cells. **C**. Spike expression in pseudovirus-producing cells stained with a polyclonal rabbit antibody (See supplementary Methods for details).

Infected or vaccinated participants are summarized in Table 1. The study was approved by the Ethics Committee Board from Hospital Universitari Germans Trias i Pujol (HUGTiP, PI-20-122 and PI-20-217) and all participants provided written informed consent before inclusion. Plasma samples were obtained from the prospective KING cohort of the HUGTiP (Badalona, Spain) and from Althaia (Manresa, Spain) from March 2020 to February 2021; thus, covering the different COVID-19 outbreaks in Catalonia (dadescovid.cat). We analyzed 32 individuals infected in March-2020, using plasma samples collected 48 days (n=16) or 196 (n=16) days after symptoms onset. We also selected 16 individuals infected in August 2020 using plasma samples collected 44 days after symptoms onset, and 5 patients (n=13 samples) infected in January 2021 by the B.1.1.7 variant. Finally, 32 individuals having received two doses of Pfizer/Biontech vaccine were sampled 2 weeks after the second dose. This last group included uninfected and long-term previously infected individuals. Description of the different groups and subgroups is shown in Table 1.

**Table 1.**
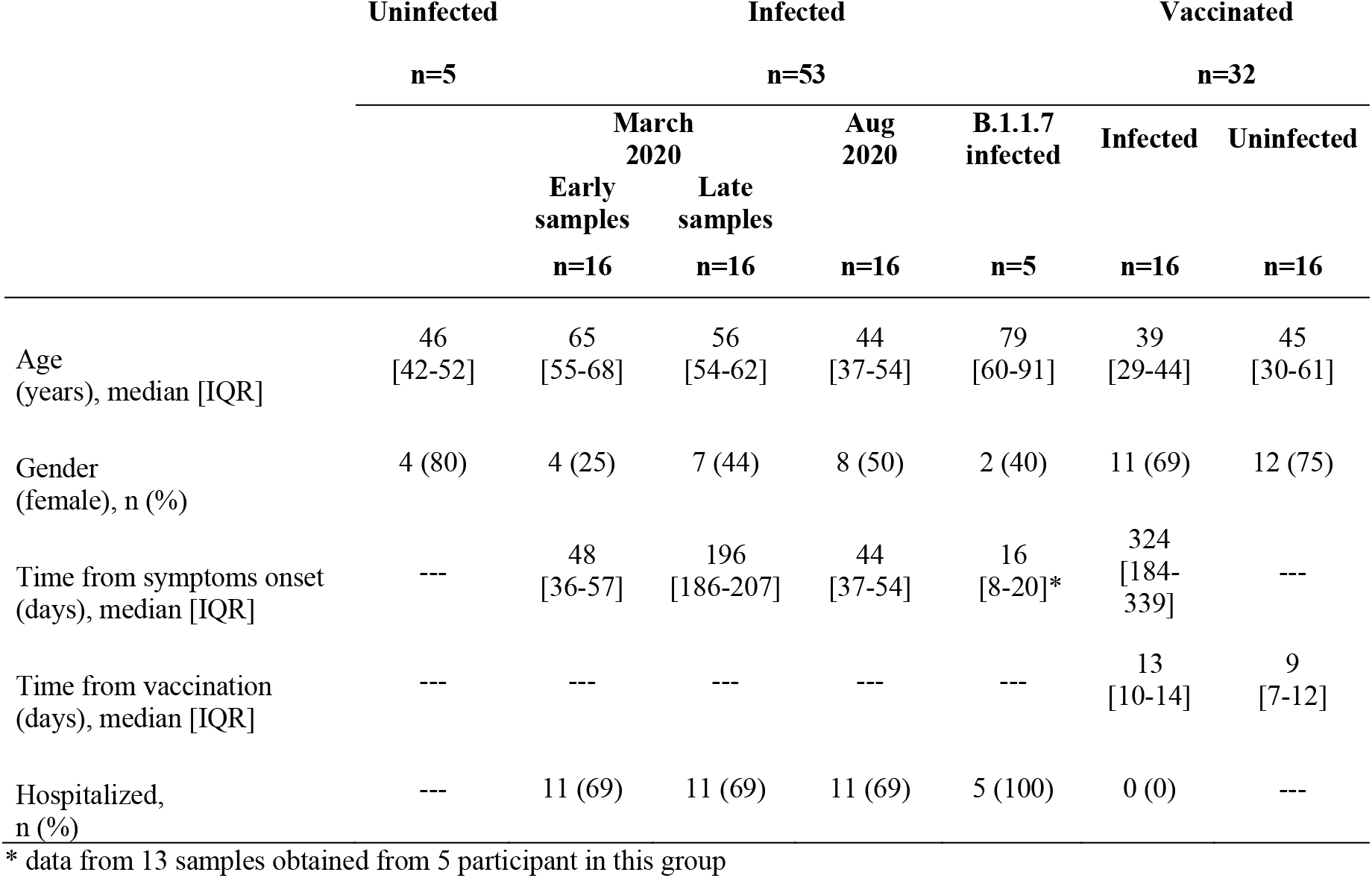
Description of participants. Uninfected individuals were included as negative controls for neutralizing activity. All of them showed undetectable neutralizing activity.

We tested all plasma samples (n=98) against the three SARS-CoV-2 pseudoviruses and analyzed the data by comparing vaccinated and non-vaccinated individuals. The neutralization titer of vaccinated individuals against the B.1.1.7 and the original Wuhan virus was not statistically different (ANOVA test with Dunn’s correction), despite Wuhan variant expressing the matched sequence of the vaccine (median change was 0.97). In contrast, vaccinated individuals showed significantly higher potency to neutralize the intermediate D614G mutant (*p*=0.001). A similar analysis including all non-vaccinated infected individuals showed similar results. The highest neutralization (*p*=0.001) was noticed for the D614G mutant, while no significant differences were observed between the Wuhan and the B.1.1.7 pseudoviruses (median fold-change of 1.44, *p*=0.112). However, this value was significantly different from the value obtained in vaccinees (M-W test, Figure 2).

**Figure 2.**
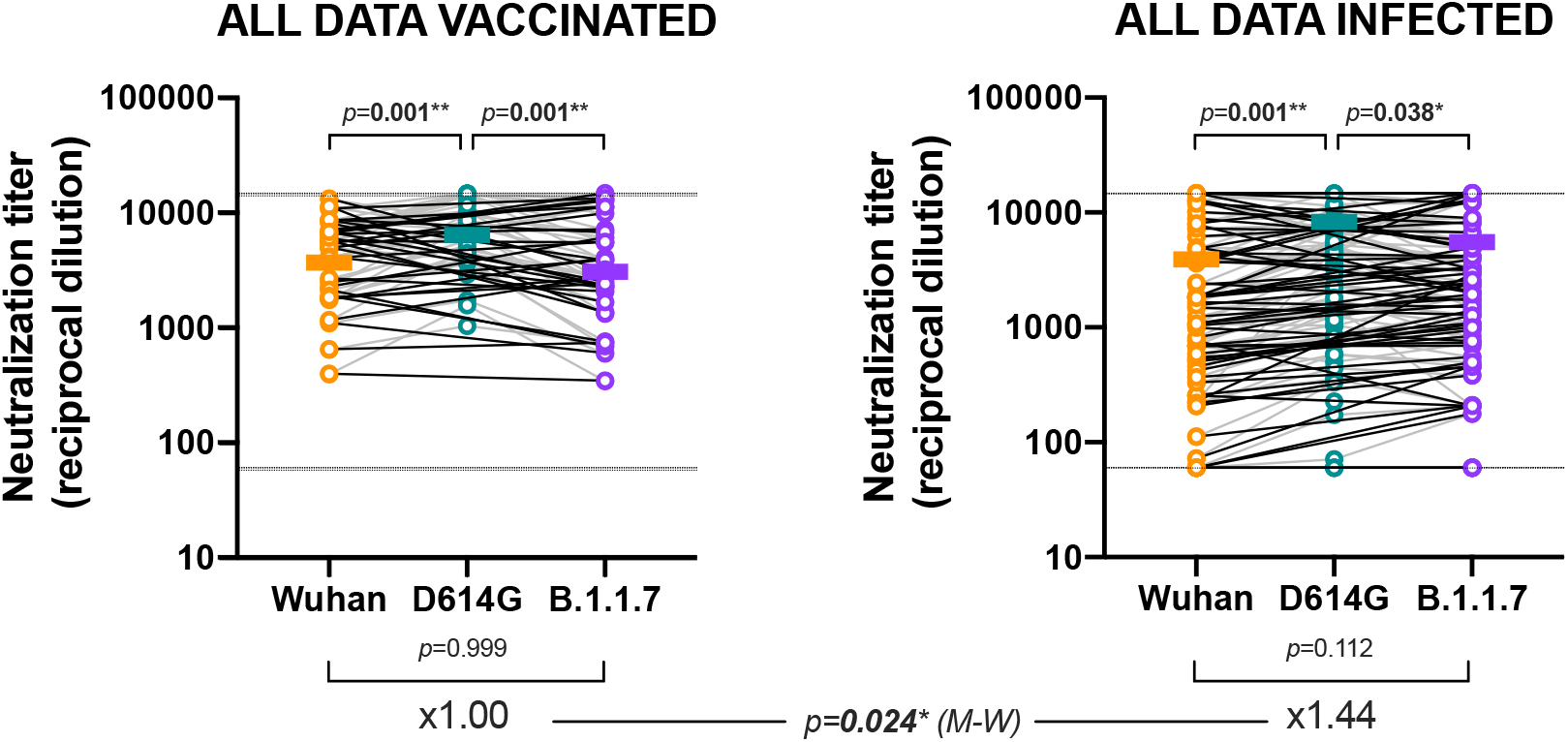
Global analysis of neutralization titers in infected and vaccinated individuals. Values of ID50 (as reciprocal dilution) are shown for all plasma samples from vaccinated and infected individuals against the indicated pseudoviruses. Top and bottom *p* values show the comparison of median titers among the three viruses (Friedman test with Dunn’s multiple comparison test). Bottom values indicate the median fold-change between Wuhan and B.1.1.7 isolates. Fold change medians were compared using Mann Whitney test (middle *p*-value).

To understand these differences, we analyzed infected and vaccinated subgroups. To assess the impact of sequence evolution on the immune responses, infected individuals were grouped according to infection date. Individuals infected during the first wave (March 2020) were exposed in majority to the D614-containing variants (60% prevalence at that time in Spain), while individuals infected during the second wave (August 2020) were largely exposed to the G614-containing 20E(EU1) lineage, which accounted for nearly 100% of new infections during the summer of 2020 (https://nextstrain.org/ncov/global). Individuals infected by the B.1.1.7 variant identified in January 2021 were also analyzed. Longer follow up of patients infected in march 2020 allowed for the analysis of long-term (>6 months) neutralizing activity (8). Vaccinated individuals were subclassified according to previous COVID-19 evidence in infected or uninfected.

Individuals infected on March 2020 sampled 48 days or >6 months after infection showed a non-significant change of neutralization activity comparing Wuhan and B.1.1.7 variants (median =1.03-fold change in early and 1.68-fold in late sampling groups). A direct comparison of these median fold-changes showed statistically significant difference between groups (p=0.036, M-W test, Figure 3A) indicating that time from infection modulates changes in cross-neutralization activity. This observation supports the positive evolution overtime of neutralizing responses in infected individuals suggested by different authors (8,10).

**Figure 3.**
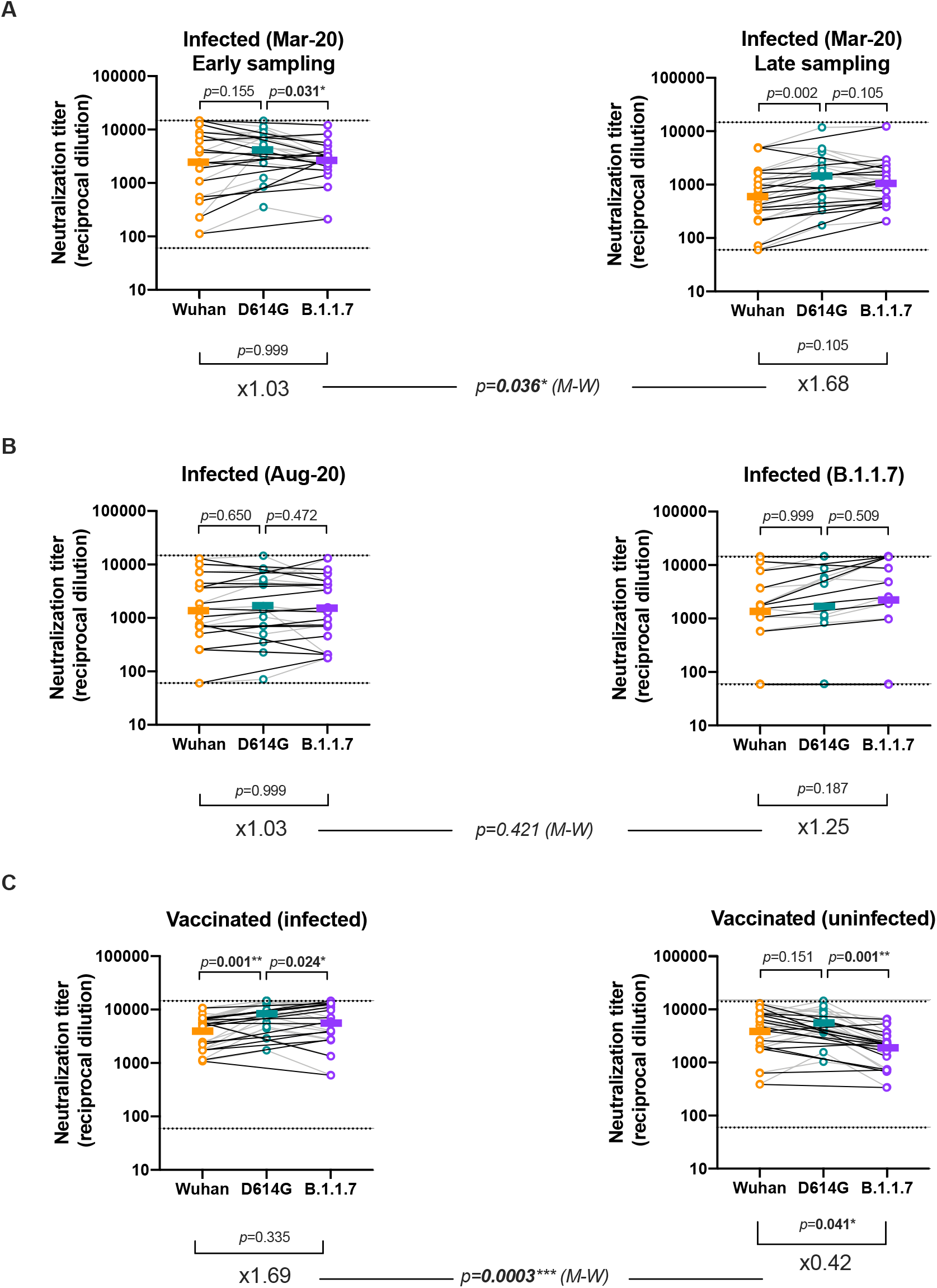
Subgroup analysis of neutralization titers in infected and vaccinated individuals. Values of ID50 (as reciprocal dilution) are shown for the indicated plasma samples against the different pseudoviruses as in Figure 2. Top and bottom *p* values show the comparison of median titers among the three viruses (Friedman test with Dunn’s multiple comparison test). Bottom values indicate the median fold-change between Wuhan and B.1.1.7 isolates. Fold change medians were compared using Mann Whitney test with false discovery rate correction (middle *p*-values).

Individuals infected in August 2020 or infected by the B.1.1.7 variant behaved differently from those infected in March 2020. No significant changes were noticed between neutralizing titers against the three tested viruses (Figure 3B). This observation is consistent with the fact that all these individuals were infected with viruses containing the G614 mutation and suggests that antibodies elicited by the 20E(EU1) and B.1.1.7 variants retain neutralizing activity against former viral sequences.

Finally, the analysis of vaccinated individuals yielded some relevant differences. Both subgroups (infected and non-infected) showed an opposite change in neutralization activity comparing Wuhan and B.1.1.7 variants (median 1.69-fold in infected and 0.42 in uninfected vaccinated), being the decrease in the uninfected groups significant (*p=*0.041). A direct comparison of ratios showed that this opposite behavior was statistically different between groups (*p*=0.0003) indicating that natural pre-infection increases the ability of antibodies elicited by vaccines to neutralize the B.1.1.7 variant (Figure 3C). This observation is consistent with data from infected non-vaccinated individuals, in which both the time from infection and the infecting virus help maintaining neutralization activity. Since the group of vaccinated individuals is heterogeneous in time from infection, a wider analysis will be necessary to dissect the contribution of each factor. In addition, our data do not allow us to draw any conclusion on the long-term evolution of immune responses elicited by vaccines, in terms of quality of antibodies or durability.

In summary, we show that increased infectivity, *in vitro,* of the B.1.1.7 variant minimally impacts sensitivity to neutralizing immune responses from infected and/or vaccinated individuals, as previously reported (4,11). Importantly, our data suggest that previous infection contributes to a better neutralization of B.1.1.7 variant in vaccinees, being the uninfected vaccinees the individuals showing a more evident, although small in magnitude, loss of neutralization (12,13). Both wider diversity and evolution of the antibody repertoire could contribute to this effect (14), which is also observed in infected non-vaccinated individuals (Figure 3A). In line with other reports (15), we demonstrate that the B.1.1.7 variant elicits cross-neutralizing responses against former viral variants, thus determining potential viral evolution.

## Acknowledgements

This work was partially funded by Grifols, the *Departament de Salut* of the *Generalitat de Catalunya* (grant SLD0016 to JB and Grant SLD015 to JC), the Spanish Health Institute *Carlos III* (Grant PI17/01518 and PI18/01332 to JC), CERCA Programme/*Generalitat de Catalunya* 2017 SGR 252, and the crowdfunding initiatives #joemcorono, BonPreu/Esclat and Correos. The funders had no role in study design, data collection and analysis, the decision to publish, or the preparation of the manuscript. EP was supported by a doctoral grant from National Agency for Research and Development of Chile (ANID): 72180406. We are grateful to all participants and the technical staff of IrsiCaixa for sample processing.

## Author Contributions

BT, JB and BC designed and coordinated the study. EP, SM, CR, FT-F, RO, JV-A, JS and NI-U performed and analyzed neutralization assays. RL, VG and VU performed structural and statistical analysis. GT, JT, CG-F, AF, RPV, AC, RP, IB, EG, MM and JC selected patients and coordinated data. JB drafted the manuscript and all authors have made substantial contributions to the revision of the subsequent versions. All authors approved the submitted version of the manuscript and agreed both to be personally accountable for the author’s own contributions and to ensure that questions related to the accuracy or integrity of any part of the work.

