## Supplementary Methods for "Previous SARS-CoV-2 infection increases B.1.1.7 cross-neutralization by vaccinated individuals"

[1]IrsiCaixa AIDS Research Institute, Germans Trias i Pujol Research Institute (IGTP), Can Ruti Campus, UAB, 08916, Badalona, Catalonia, Spain
[2]IRTA Centre de Recerca en Sanitat Animal (CReSA, IRTA-UAB), Campus de la UAB, 08193, Bellaterra, Catalonia, Spain
[3]UAB, Centre de Recerca en Sanitat Animal (IRTA-UAB), Campus de la UAB, 08193, Bellaterra, Catalonia, Spain
[4]Departament de Sanitat i Anatomia Animals, Facultat de Veterinària, UAB, 08193, Bellaterra, Catalonia, Spain
[5]Barcelona Supercomputing Center, 08034, Barcelona, Catalonia, Spain
[6]Catalan Institution for Research and Advanced Studies (ICREA), 08010, Barcelona, Catalonia, Spain
[7]Fundació Althaia, Hospital de Sant Joan de Déu, 08243, Manresa, Catalonia, Spain
[8]Infectious Diseases Department, Fight against AIDS Foundation (FLS), Germans Trias i Pujol Hospital, 08916, Badalona, Catalonia, Spain
[9]Germans Trias i Pujol Hospital, 08916, Badalona, Catalonia, Spain
[10]University of Vic–Central University of Catalonia (UVic-UCC), 08500, Vic, Catalonia, Spain
[¶]Equal contribution

***Corresponding author:**

Julià Blanco, PhD
Senior Researcher
Institut de Recerca de la Sida. IrsiCaixa, IGTP, Hospital Germans Trias i Pujol
Ctra. de Canyet s/n. 2a Planta Maternal. 08916 Badalona. Barcelona


### SUPPLEMENTARY METHODS

### EXPERIMENTAL MODEL AND SUBJECT DETAILS

#### Study overview and subjects

The study was approved by the Hospital Ethics Committee Board from Hospital Universitari Germans Trias i Pujol (PI-20-122 and PI-20-217) and all participants provided written informed consent before inclusion.

Plasma samples were obtained from individuals of the prospective KING cohort of the HUGTiP (Badalona, Spain). The recruitment period lasted from March to October 2020, thus covering the first and second waves of COVID-19 outbreak in Catalonia (dadescovid.cat). The KING cohort included individuals with a documented positive RT-qPCR result from nasopharyngeal swab and/or a positive serological diagnostic test.

#### Cell lines

HEK293T cells (presumably of female origin) overexpressing WT human ACE-2 (Integral Molecular, USA) were used as target for SARS-CoV-2 spike expressing pseudovirus infection. Cells were maintained in T75 flasks with Dulbecco′s Modified Eagle′s Medium (DMEM) supplemented with 10% FBS and 1µg/mL of Puromycin.

### METHOD DETAILS

#### Spike plasmid generation

SARS-CoV-2.SctΔ19 Wuhan and B.1.1.7 were generated (Geneart) from the full protein sequence of the original Wuhan and the UK variant (B.1.1.7) spike sequences, with a deletion of the last 19 amino acids in C-terminal (1), human-codon optimized and inserted into pcDNA3.1(+). The G614 spike mutant was generated by site-directed mutagenesis as previously described (2). In brief, SARS-COV-2.SctΔ19 Wuhan plasmid was amplified by PCR with Phusion polymerase (Thermo Scientific, F-549S) and the following primers: 5'-TACCAGGgCGTGAACTGTACCGAAGTGCC-3' and 5'-GTTCACGcCCTGGTACAGCACTGCCAC-3'. PCR was 20 cycles with an annealing temperature of 60ºC and an elongation temperature of 72ºC. PCR product was then treated for 3 hours with the DpnI restriction enzyme (Thermo Scientific, ER1705), to eliminate template DNA, and transformed into supercompetent E coli. Final mutated DNA was then fully sequenced for validation.

**Pseudovirus generation and neutralization assay**

HIV reporter pseudoviruses expressing SARS-CoV-2 S protein and Luciferase were generated using the defective HIV plasmid pNL4-3.Luc.R-.E- was obtained from the NIH AIDS Reagent Program(3). Expi293F cells were transfected using ExpiFectamine293 Reagent (Thermo Fisher Scientific) with pNL4-3.Luc.R-.E- and SARS-CoV-2.SctΔ19 (Wuhan, G614 or B.1.1.7), at a 8:1 ratio, respectively. Control pseudoviruses were obtained by replacing the S protein expression plasmid with a VSV-G protein expression plasmid as reported(4). Supernatants were harvested 48 hours after transfection, filtered at 0.45 μm, frozen, and titrated on HEK293T cells overexpressing WT human ACE-2 (Integral Molecular, USA). This neutralization assay has been previously validated in a large subset of samples(5).

Neutralization assays were performed in duplicate. Briefly, in Nunc 96-well cell culture plates (Thermo Fisher Scientific), 200 $TCID_{50}$ of pseudovirus were preincubated with three-fold serial dilutions (1/60–1/14,580) of heat-inactivated plasma samples for 1 hour at 37ºC. Then, $2x10^4$ HEK293T/hACE2 cells treated with DEAE-Dextran (Sigma-

Aldrich) were added. Results were read after 48 hours using the EnSight Multimode Plate Reader and BriteLite Plus Luciferase reagent (PerkinElmer, USA). The values were normalized, and the $ID_{50}$ (the reciprocal dilution inhibiting 50% of the infection) was calculated by plotting and fitting the log of plasma dilution vs. response to a 4-parameters equation in Prism 8.4.3 (GraphPad Software, USA).

**Flow cytometry**

Transfected Expi293 T cells were first stained extracellularly with a polyclonal rabbit anti-spike RBD antibody (Sino Biological, 40592-T62) and a secondary APC labeled anti-rabbit antibody (Jackson ImmunoResearch, 109-136-098). Cells were then fixed (Life Technologies, GAS001S100) and stained, in permeabilization buffer (Life Technologies, GAS002S100), with a FITC-labeled mouse anti-p24Gag antibody KC57 (Beckman Coulter, 6604665). Cells were acquired on a BD FACS Celesta flow cytometer using FACSDIVA 8.0.1.1 software and analyzed with FlowJo 10.62 (Tree Star, Inc, USA).

### STATISTICAL ANALYSIS

Continuous variables were described using medians and the interquartile range (IQR, defined by the $25^{th}$ and $75^{th}$ percentiles), whereas categorical factors were reported as percentages over available data. Quantitative variables were compared using the Mann-Whitney test, and percentages using the chi-squared test. Friedman test with Dunn's multiple comparison test was used to compare neutralization of different pseudoviruses. Multiple M-W comparisons were corrected by false discovery rate. Analyses were performed with Prism 8.4.3 (GraphPad Software) and R version 4.0 (R Foundation for Statistical Computing).

### Supplmentary references
